# Private Information Leakage from Polygenic Risk Scores

**DOI:** 10.64898/2026.02.16.706191

**Authors:** Kirill Nikitin, Gamze Gürsoy

## Abstract

Polygenic Risk Scores (PRSs) estimate the likelihood of individuals to develop complex diseases based on their genetic variations. While their use in clinical practice and direct-to-consumer genetic testing is growing, the privacy implications of publicly sharing PRS values are often underestimated. In this work, we demonstrate that PRSs can be exploited to recover genotypes and to de-anonymize individuals. We describe how to reconstruct a portion of an individual’s genome from a single PRS value by using dynamic programming and population-based likelihood estimation, which we experimentally demonstrate on PRS panels of up 50 variants. We highlight the risks of combining multiple, even larger-panel PRSs to improve genotype-recovery accuracy, which can lead to the re-identification of individuals or their relatives in genomic databases or to the prediction of additional health risks, not originally associated with the disclosed PRSs. We then develop an analytical framework to assess the privacy risk of releasing individual PRS values and provide a potential solution for sharing PRS models without decreasing their utility. Our tool and instructions to reproduce our calculations can be found at https://github.com/G2Lab/prs-privacy.

## 1 Introduction

A polygenic risk score (PRS) is a predictive measure that quantifies an individual’s genetic susceptibility to complex traits, based on the cumulative effect of their inherited genetic variants. PRSs have demonstrated robust predictive performance across a wide range of complex diseases [1, 2], including successful applications in clinical cohorts [3, 4, 5, 6, 7, 8]. Their utility for risk stratification has spurred discussions about integrating PRS-based assessments into routine healthcare [4, 9, 10]. Leading direct-to-consumer genetic testing companies, such as 23andMe [11] and Nebula Genomics [12], already employ PRSs to provide consumers with personalized trait risk estimates. The use of PRSs in therapeutic decision-making, disease screening, and personalized life planning is anticipated to significantly expand in the coming years [10].

Whereas ethical and policy debates on the commercial use of PRSs are beginning to take shape [13], the privacy implications of sharing individual scores remain largely unexplored. PRSs are commonly viewed as summary-level data due to the assumption that a single value lacks sufficient granularity to disclose any additional information, beyond the assessed trait’s risk. As a result, genetic association studies periodically publicly release individual PRSs [14, 15, 16], and customers of direct-to-consumer genetic testing companies share their scores on online forums to seek health advice and recommendations [17, 18, 19]. PRSs are, however, derived from genetic data, which have long been regarded as highly sensitive [20, 21], due to their potential to re-identify individuals [22, 23, 24, 25, 26] or to reveal their health status [27, 28]. As the use of PRSs becomes increasingly integrated into clinical practice and research, a deeper understanding of the privacy implications associated with sharing this information is essential.

In this study, we present the first investigation into the potential leakage of sensitive genetic information through PRSs and evaluate scenarios in which this poses significant privacy risks. By framing genotype recovery from PRS values as the subset-sum problem, we have developed an efficient dynamic-programming algorithm that uses basic population genetic statistics to accurately infer the underlying genotypes from PRS values. We find that the risk of disclosing patient-level PRS values depends on the parameters of the PRS models, specifically the number of loci and the precision of the effect weights, and that models with up 80 variants are vulnerable to genotype recovery in practice. Based on this, we first analyze 4,723 PRS models published in the PGS Catalog [29] and find that at least 447 of them would be vulnerable if patient-level PRSs were shared. We then take a subset of the vulnerable models of up to 50 variants each and experimentally demonstrate that given a panel of PRSs, e.g., from a direct-to-consumer genetic-testing report [11], an attacker can reconstruct the associated genotypes of an individual with 95% accuracy. Notably, individuals of non-European ancestry appear to be at a greater leakage risk due to biases in existing Genome-Wide Association Studies (GWAS). The recovery is then sufficient for the re-identification of the individuals or their genetic relatives in genealogy databases with near 100% accuracy. We also demonstrate that a typical published PRS based on just 27 loci is sufficient to uniquely identify 95% of individuals in large cohorts with 450K participants. Finally, we develop an analytical framework to assess the privacy risks associated with sharing PRS values and models, and propose a simple yet effective solution that preserves individual privacy while maintaining the utility of shared information.

We analyze three distinct privacy threat models related to PRSs, each with different adversarial capabilities and access assumptions. First, we study genotype recovery from published PRSs under the assumption that model parameters are publicly available (§2.2). Second, we examine genealogy-based re-identification that uses recovered genotypes and public genetic genealogy databases (§2.3). Third, we evaluate PRS-based linkage risk under an explicit access model in which an adversary can compute PRSs for individuals in an anonymized genotype-phenotype database (§2.4). These scenarios represent qualitatively different attack settings and are analyzed separately throughout the manuscript.

## 2 Results

### 2.1 PRS-based genetic privacy attacks

A polygenic risk score (PRS) is a numerical value that quantifies an individual’s genetic predisposition to a particular disease or trait. It is calculated by aggregating the effects of multiple genetic variants. A PRS is by definition a summary statistic derived from a patient’s genetic data, but it is generally considered not to contain any directly identifiable private information about the patient. In this study, we explored whether private genetic variants could be recovered from PRSs and whether these variants could then be used to infer additional sensitive information about patients.

Our attack framework (Fig. 1) is based on two key scenarios: (1) If a research study publicly releases anonymized individual PRSs [14, Supplementary Table 5][15, Figure 1], or an anonymous user discloses their PRS online to seek health advice [17, 18, 19], it can be possible to infer the associated genotypes of these anonymized patients and to cross-reference genealogy websites to re-identify the patient or their relatives. Note that this is achieved by *querying* genealogy websites, as they typically restrict users from downloading participants’ genomic data. (2) If a known patient publicly shares their PRS (e.g., through social media or online forums, such as WebMD), it can be possible to infer their associated genotypes and to search anonymized genotype-phenotype databases with restricted access (e.g., genomic beacons [30] or any database that permits only query-based access) to uncover more sensitive phenotypic information. In both scenarios, further analysis could reveal the predisposition to diseases that the original PRS did not include.

**Fig. 1.**
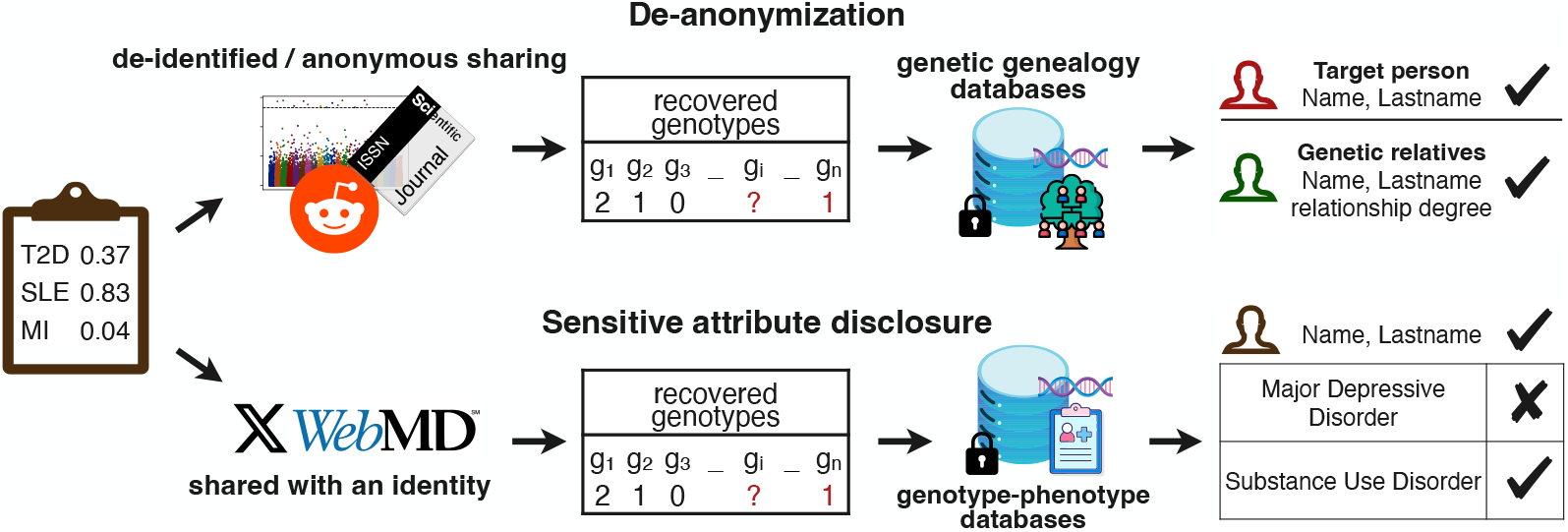
Overview of privacy risks with sharing PRSs. This illustration depicts the potential privacy risks linked to predicting SNP genotypes from PRS values. For publicly released PRS values of anonymous individuals or from anonymous GWAS participants, the predicted genotypes can be used to de-anonymize them or their genetic relatives by cross-referencing genetic genealogy databases. In cases where PRS values are released for known individuals, the predicted genotypes can be used to infer sensitive health information by searching genotype-phenotype databases.

If a profit-driven entity, such as an insurance provider, gains access to the PRS values and genetic ancestry of a target individual, as well as the corresponding effect weights and SNP locations, both of which can be obtained from resources, such as the Polygenic Score (PGS) Catalog [29], it could reconstruct the genotypes for the SNPs included in the PRS panel. These reconstructed genotypes can then be used to: (a) identify the target individual or their genetic relatives in genealogy databases, (b) infer other potential diseases that the target might have by cross-referencing query-only genotype-phenotype databases, or predict predisposition for additional diseases of the target individual. In all cases, sensitive information about the target is exposed, either revealing their identity, if the PRSs are anonymized, or disclosing private health details, if their identity is already known. Although the Genetic Information Nondiscrimination Act (GINA) [31] protects from genetic-risk discrimination in health insurance, it does not extend to disability, property, or life insurances.

Our goal is to quantify the extent to which an individual’s PRS reveals sensitive information, identify the conditions that amplify or mitigate this exposure, and develop strategies to reduce associated privacy risks. To achieve this, we developed methods that leveraged publicly available data—such as lists of PRSs, genomic locations of SNPs included in the PRS models, and their corresponding effect weights—to infer an individual’s associated SNP genotypes. A PRS for a disease is the linear combination of the genotypes of the associated SNPs, defined as *PRS* = ∑*β*_*j*_ · *g*_*j*_, where *β*_*j*_ is the effect weight of SNP *j* and *g*_*j*_ is the genotype of SNP *j* (0 for homozygous reference alleles, 1 for heterozygous alternative allele, and 2 for homozygous alternative alleles). Inferring the genotypes from a PRS value and effect weights can be framed as a variation of the subset-sum problem [32]. The subset-sum problem is a computational problem where, given a set of numbers, the goal is to find a subset whose sum equals a target value. It is known to be NP-hard [33], meaning that it can be computationally difficult to solve, especially as the set size grows. Here, we reduce the genotype recovery task to an instance of the subset-sum problem by defining the effect weights as the number set and the PRS value as the target sum. The number of times the solution contains each effect weight (0, 1, or 2) determines the genotype for each corresponding SNP.

Our framework for inferring a target individual’s SNP genotypes based on a PRS value and the associated effect weights consists of several steps. First, it assesses the solvability of the problem by using the concept of *density* [34]. In subset-sum problems, density represents the ratio between the size of the set (i.e., the number of weights in our case) and the length of the bit representation of the largest weight. Higher density indicates that there are many weights relative to the bit length of the largest weight, which leads to a tighter distribution of possible subset sums. This dense spacing increases the number of combinations that could potentially match the target sum, thus making the problem more challenging to solve [35]. We adapted this concept to genetic data by applying a density threshold to determine whether a PRS is solvable—that is, whether we can expect to accurately infer the genotypes for all the associated SNPs of a target individual, given the effect weights and the PRS value (see Methods). Naturally, a PRS with fewer associated SNPs or more precise weights are easier to solve, as the computational problem becomes sparser and more tractable.

Once the PRS is determined to be solvable, we proceed to infer the associated SNP genotypes. To achieve this, we developed a dynamic programming algorithm [36], enhanced by the meet-in-the-middle approach and further optimized by using additional techniques [37] (Fig. 2A). This algorithm efficiently identifies all valid solutions, i.e., all possible genotype sets that result in the target PRS. In order to assess the plausibility of each potential solution, we calculate a log-likelihood score by using allele frequencies from the target individual’s population. Specifically, we compute the sum of the log-probabilities for the identified genotypes in each solution. The core idea is that a solution that closely aligns with the population average is more likely to be correct. The solution with the highest total sum, which corresponds to the highest likelihood, is then selected as the most probable genotype configuration (Fig. 2A). We then incorporate continuous log-likelihood estimation directly into the dynamic-programming algorithm, which enables us to handle PRSs with more SNPs. Additionally, we implement two techniques to further increase the solving accuracy and the size of solvable PRSs that we call PRS chaining and self-repair. In PRS chaining, we first solve the PRS with the smallest set of SNPs. For each subsequent PRS that includes more SNPs, we retain the genotypes of overlapping SNPs from the previous solution. This approach gradually reduces the possible solution space as the number of SNPs increases, thereby making the computations more efficient (Fig. 2B). When incorporating solutions from a previous PRS, if the solution for the current PRS fails, it suggests that the integrated genotypes might be incorrect. This enables us to revisit and to correct these genotypes, a process that we refer to as *self-repair*. For details, see Methods.

**Fig. 2.**
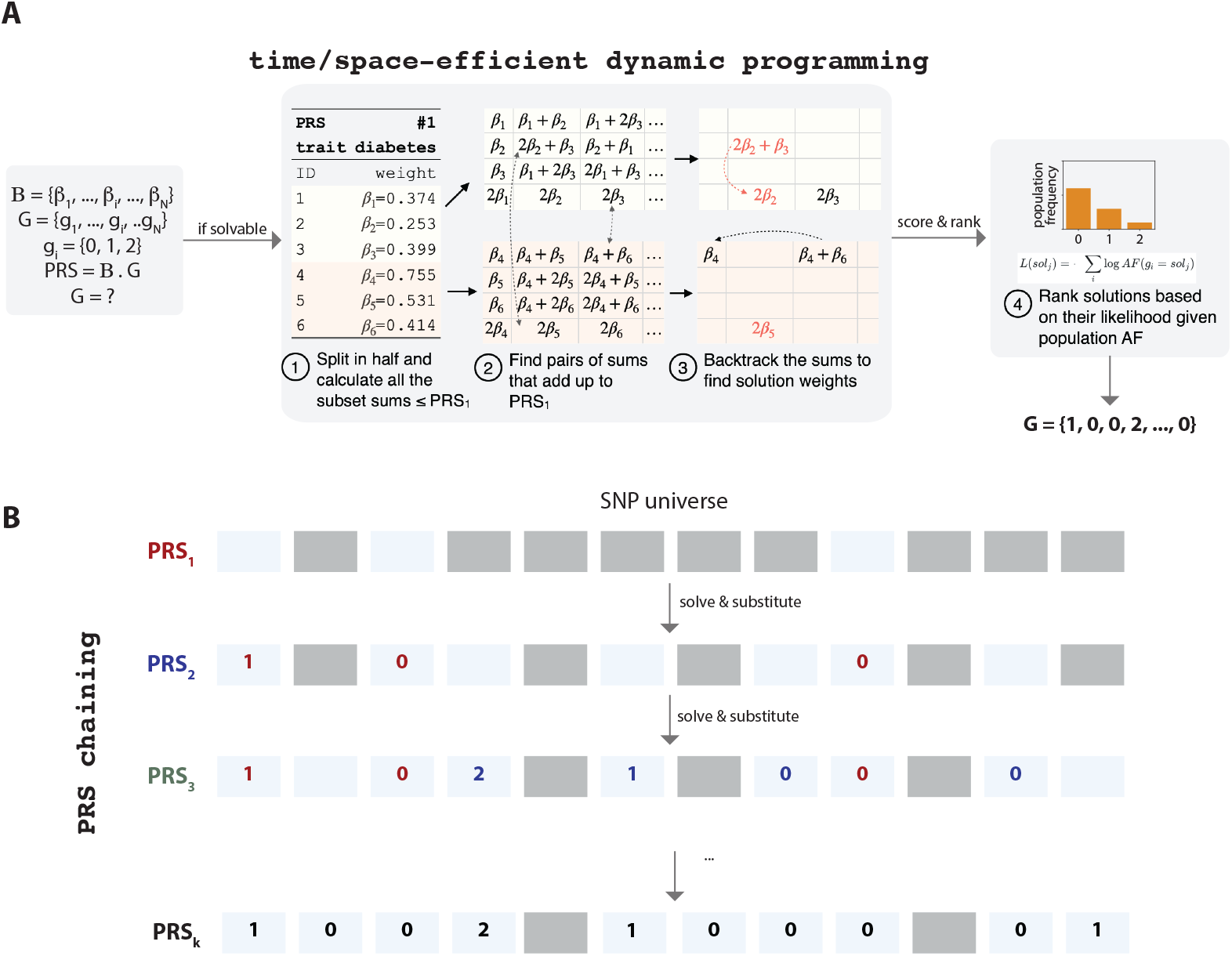
Framework for predicting genotypes from PRS values and PRS chaining. **(A)** Illustration of the genotype prediction process via dynamic programming. Given a set of effect weights and a target PRS, the goal is to find the unknown genotypes. We first determine the solvability of a PRS through density calculation. If the PRS is solvable, the framework uses a dynamic programming algorithm to find all possible solutions by using a meet-in-the-middle approach. The meet-in-the-middle approach divides the effect weights into two parts, creating separate tables of partial sums for each. It then identifies pairs of entries from these tables that sum to the target PRS value. Partial sums are added only if the target remains achievable. The search is further refined in two stages: first, rounded-down weights identify solution candidates, and then, precise matches to the target are selected. These strategies significantly reduce the memory and computational costs, which facilitates the genotype prediction from PRSs with a larger number of variants. We then score and rank potential genotype solutions based on population allele frequencies. **(B)** Schematic of our PRS-chaining technique. Starting with a PRS involving a small set of SNPs, the predicted genotypes for overlapping SNPs are substituted into progressively larger PRSs. This iterative process narrows the solution space and thus increases the efficiency of genotype prediction as more SNPs are incorporated across multiple PRSs.

### 2.2 Patient genotypes can be recovered from publicly available PRSs

The solvability of the genotype recovery problem is determined by the parameters of the corresponding PRS model, specifically the number of associated SNPs and the precision (i.e., the number of decimal digits) of the effect weights. By following by the literature on the subset-sum problem and via experimental evaluation, we determined the range of parameters where genotype recovery was feasible (see Methods). We then analyzed all PRS models published in the PGS Catalog [29] to assess the proportion that could be resolved if individual PRSs were available. We found that at least 447 out of 4,723 published PRS models were vulnerable to genotype recovery.

Correctly inferring genotypes from a single PRS might already suffices for de-identifying an individual, as prior research demonstrated that as few as 20 SNP genotypes could enable re-identification [22, 23]. In practice, an attacker can obtain a list of PRS values for a single individual, e.g., an insurance company might receive a patient’s genetic-testing report with over 40 risk scores [11]. Combining prediction results from multiple PRSs, even those with more SNPs than theoretically solvable, enables an attacker to achieve substantial coverage of an individual’s genome.

We evaluated the genotype recovery ability of the attacker by using whole-genome sequencing data from the 1000 Genomes Project (2,535 samples in total, 50 individuals with first-degree and 18 individuals with second-degree relationships [38]) and a PRS set containing scores of up to 50 SNPs each from the PGS catalog for various diseases. After filtering the PRSs based on our solvability threshold and performing quality controls (see Methods), we selected a set of 298 PRSs, encompassing 4,821 SNPs, of which 2,654 were unique (Supplementary Table 1). We then predicted the genotypes of these SNPs for each individual given their PRS values. We compared the results against two baselines: the deterministic predictor that assigned the major genotype of the individual’s ancestry at each SNP, and the stochastic predictor that sampled genotypes according to ancestry-specific probabilities.

Our method achieved a median genotype prediction accuracy of 94.6% (Fig. 3A), significantly outperforming both baseline approaches. On average, we correctly predicted 2,450 genotypes per individual. Notably, we observed higher prediction accuracy for individuals of African (AFR) and East Asian (EAS) genetic ancestries compared to those of European (EUR) ancestry. We hypothesized that this increased accuracy resulted from a higher proportion of loci where the effect allele frequencies (EAFs) were close to 0 or 1 in the AFR and EAS populations.

**Fig. 3.**
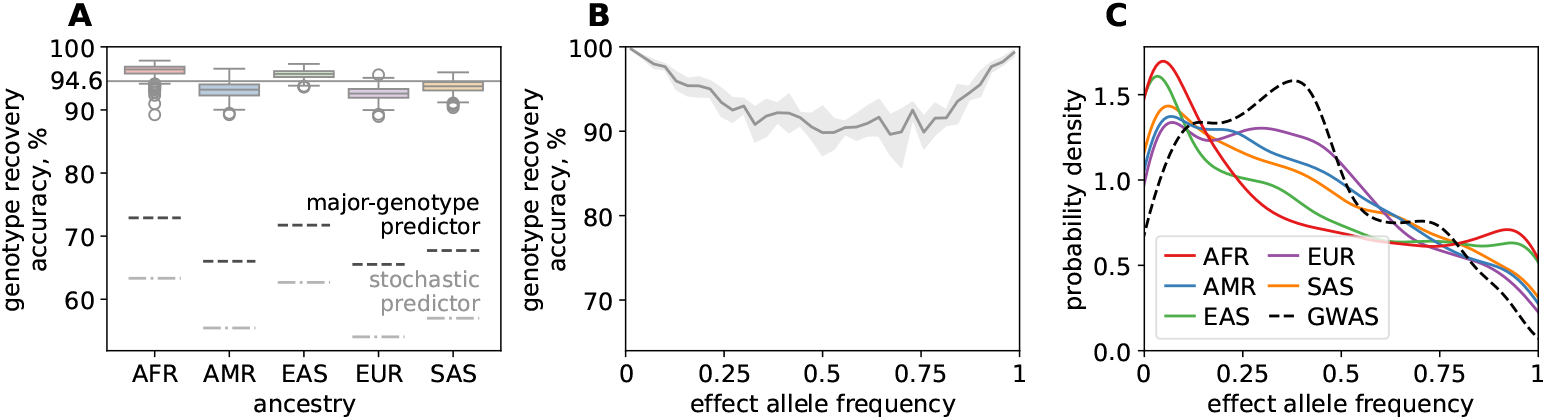
Genotype recovery accuracy. **(A)** Genotype recovery accuracy across different populations: African (AFR), Admixed American (AMR), East Asian (EAS), European (EUR), and South Asian (SAS). The baselines represent the recovery accuracy of predicting the major genotype or stochastically sampling a genotype for each SNP for every individual, according to the population allele frequencies, without access to the PRS values. **(B)** The relationship between genotype recovery accuracy and effect allele frequency. It is easier to predict genotypes for SNPs whose effect allele or other allele frequency is rare. **(C)** The distribution of effect allele frequencies in different populations compared to those in GWASs from which the PRSs are derived from. Since individuals of EUR genetic ancestry dominate most GWAS datasets, the effect allele frequency distribution in GWASs closely mirrors that of the EUR population.

To test this hypothesis, we analyzed genotype recovery as a function of population EAF and confirmed that recovery was more challenging for loci where the EAF was close to 0.5 (Fig. 3B). At these loci, both alleles were nearly equally frequent, which led to higher heterozygosity and made accurate genotype inference more difficult. In contrast, when the EAF was near 0 or 1, one allele predominated in the population, resulting in a higher proportion of individuals being homozygous at these loci. This reduced genetic variation increased the likelihood of correctly inferring an individual’s genotype since they were more likely to carry the predominant allele.

We investigated why individuals of the AFR and EAS ancestries tended to carry more effect alleles at the loci where the allele frequencies in their populations were close to 0 or 1. We found that this phenomenon was primarily due to the over-representation of PRSs derived from GWASs that were conducted predominantly on the EUR ancestry cohorts (Fig. 3C), which confirmed the findings of prior studies on disparities in clinical studies [39]. The allele frequencies in the EUR population resembled their effect allele frequency in the GWASs. This bias led to significant discrepancies between the effect allele frequencies reported in the GWASs and the actual allele frequencies in the AFR and EAS populations. As a result, individuals from these populations were more susceptible to genotype recovery attacks because their genotypes at these loci were more predictable.

We then asked the question of whether the high genotype recovery accuracy depended on the assumption that the attacker knew the target individual’s genetic ancestry. To test this, we selected ten individuals from each ancestry group. We then performed genotype recovery with our dynamic-programming approach and scored the solution with the likelihood estimation using a universal, ancestry-agnostic allele frequency. The median accuracy across all fifty individuals was 92.7%, a slight decrease from the accuracy obtained with ancestry-specific data (Supplementary Fig. 2). As expected, individuals of the AFR and EAS ancestries showed the largest drop in accuracy, with a decrease of 3%. These results indicated that, while precise population statistics enhanced genotype recovery accuracy, our method remained robust even when the attacker lacked the knowledge of the target individual’s specific ancestry.

### 2.3 Patients and their relatives can be identified in genealogy databases by using PRSs

If PRSs are publicly shared for anonymized study participants [16] or posted online by anonymous users seeking health advice [17, 19], there is a potential risk that an attacker could re-identify the individuals. To test this risk, we designed an attack strategy that involved querying a genetic genealogy database, such as GEDMatch [40], with the genotypes inferred from the published PRSs. This approach could potentially reveal the identity of the target individual or their close relatives. It is important to note that genealogy databases do not provide direct access to individuals’ genomes; instead, they allow users to query the database to identify genetic relatives.

Although genealogy services do not publicly disclose the specific algorithms that they use for identifying relatives, prior studies [41, 42] suggest that these services search for uninterrupted genomic segments with no mismatching homozygous sites between pairs of individuals. To assess whether the predicted genotypes from our genotype recovery process were sufficient for de-anonymization, we conducted a relative search process by using the 1000 Genomes dataset as the genealogy database. This database contains genomes from 2,467 unrelated and 68 related individuals. We also used the KING-robust algorithm [43] as the relatedness inference method. It uses a large set of autosomal SNPs to calculate kinship coefficients, which help identify relationships, such as first- or second-degree relatives. The exact number of SNPs that it uses can vary depending on the dataset, but having a large number of SNPs enhances the accuracy of kinship coefficient estimates.

Despite our genotype predictions being based solely on SNPs inferred from the PRSs, we demonstrated that they were sufficient to achieve 100% precision and recall in identifying individuals, i.e., our test correctly identified all the 2,535 individuals as themselves. We also achieved ∼90% precision and recall in determining first-degree relatives, and ∼85% precision and ∼75% recall in determining second-degree relatives. This was notable compared to the 0% accuracy, achieved when using a baseline approach that simply predicted the major genotype for all the associated SNPs (Fig. 4B). Importantly, while our genotype predictions were not perfectly accurate and covered only about half the SNPs of what was considered sufficient for relatedness analysis, they still provided enough information to distinguish different levels of relatedness. The matching precision was comparable between individuals of all ancestries, which implied that a higher genotype prediction accuracy did not always translate into a higher risk of re-identifiability. That is, correctly predicted SNPs that are common within a population contribute little to individual distinctiveness. Kinship coefficients are an absolute metric, hence their performance does not directly depend on the size of the database. We demonstrate the distribution of the KING kinship coefficients from our analysis in Fig. 3A that indicates a clear separation between self-identification and unrelated individuals or even first-degree relatives. This mirrors the relationship between relatedness and genotype density observed in the original KING study [43]. Additionally, we replicated this analysis using GCTA (a statistical method for heritability estimation [44]) and found that, while GCTA exhibited slightly lower accuracy, it still successfully identified individuals and most of their relatives as related (Supplementary Fig. 3B).

**Fig. 4.**
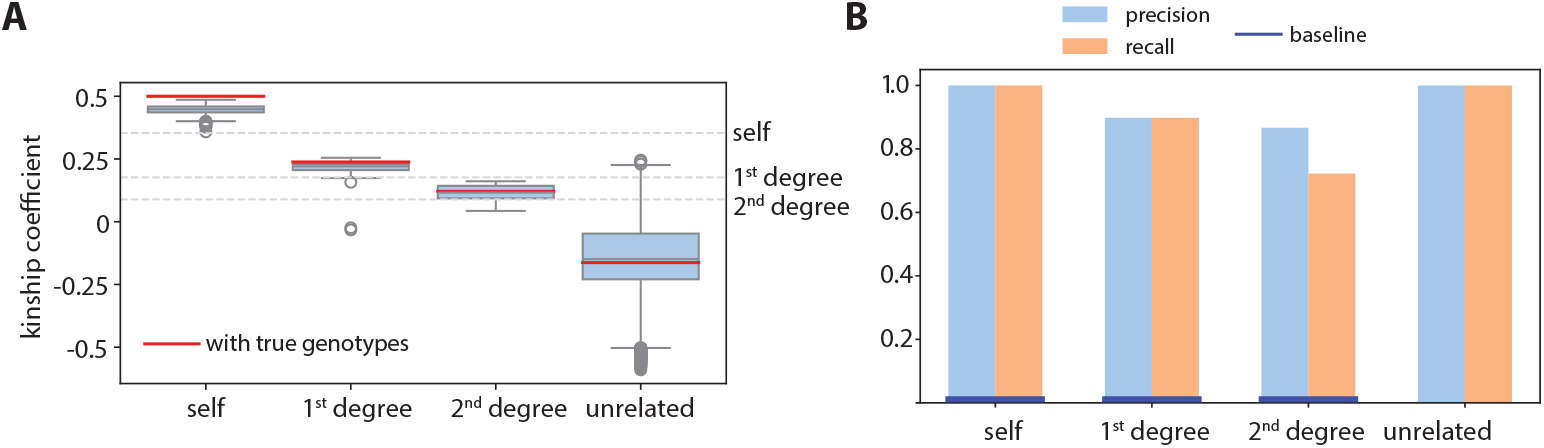
Comparison of kinship coefficient estimations and relationship classification performance. **(A)** Distribution of kinship coefficients for different relationship categories (self, 1^st^ degree, 2^nd^ degree, and unrelated) calculated using the predicted genotypes in the KING-robust. The red lines are calculated using the correct genotypes for the same SNPs. **(B)** Precision and recall metrics for identifying self, 1^st^ degree, 2^nd^ degree relationships using the predicted genotypes in the 1000 Genomes Database. Baseline performance reflects the re-identification approach of predicting the major alleles for the SNP genotypes.

### 2.4 Anonymized genotype databases can be de-anonymized by using a single PRS from known patients without the need for genotype prediction

Previous studies have shown that even a few dozen SNPs can uniquely identify an individual [22, 23]. Building on this, we examined whether a single PRS could distinguish an individual within a large genotype-phenotype database. In this scenario, we assume direct access to an anonymized genotype-phenotype database, such as UK Biobank, to which insurance companies are known to have access [45, 46]. We tested if, given an anonymized genetic database, the PRS of a known individual for a particular trait or disease, and the effect weights of the SNPs in the PRS model, it would be possible to calculate PRS values for all individuals in the database and identify the matching one (Fig. 5A). This approach relies on determining how unique PRS values are across individuals, raising the possibility that an attacker could use a single PRS to link a known individual to an anonymized dataset and to potentially expose sensitive genetic and phenotypic information. The viability of this method depends on whether the PRS for a given trait or disease is sufficiently unique to match to a specific individual in the dataset.

**Fig. 5.**
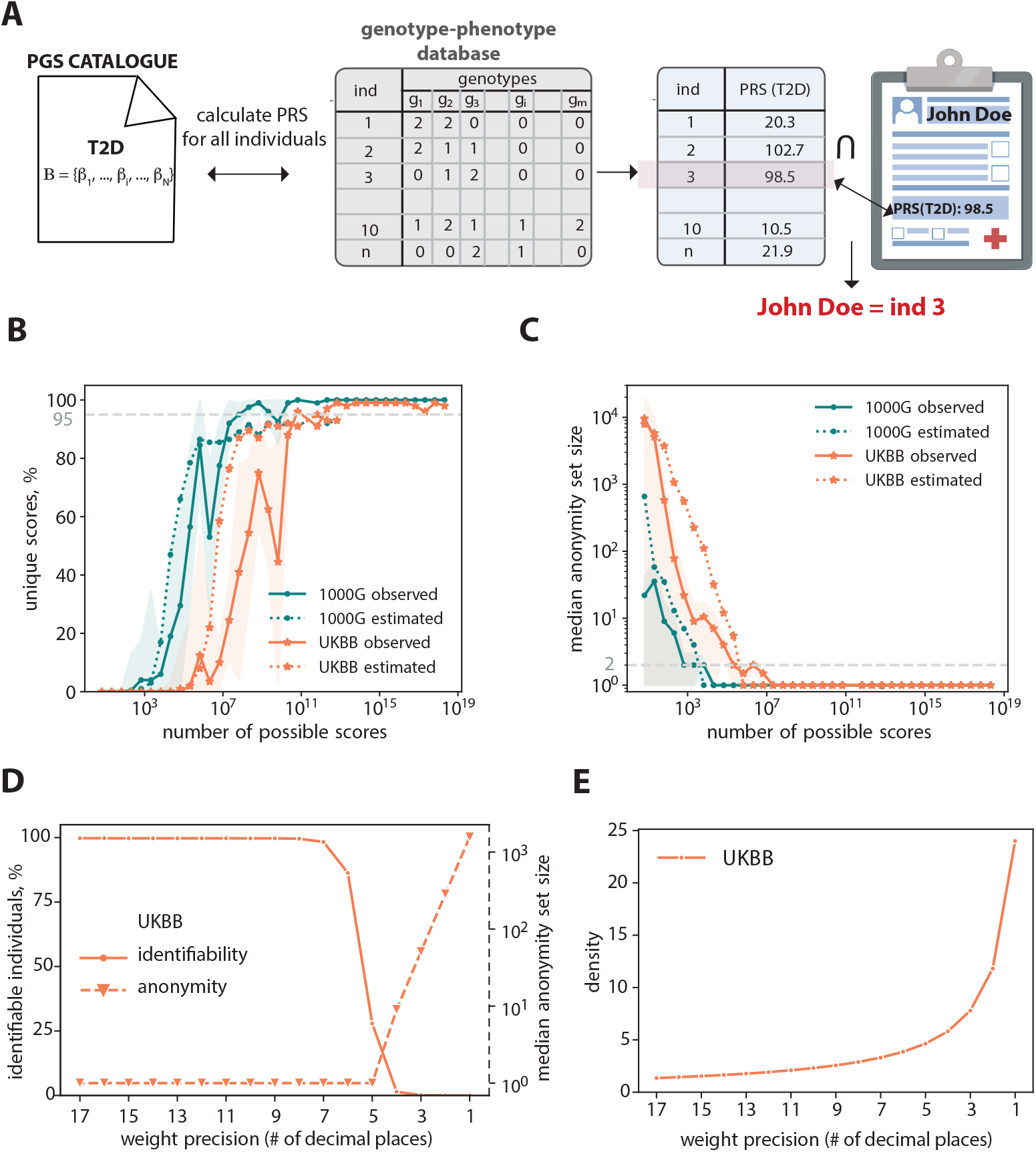
Uniqueness of PRS values. **(A)** Linking released PRS values to an anonymized genotype-phenotype database by taking advantage of the uniqueness of the PRS values. **(B)** Percentage of unique PRS values as a function of number of possible scores both observed and estimated using 1000 Genome and UKBB datasets. We found that 95% uniqueness achieved in UKBB with ∼10^11^ possible scores, which corresponds to a PRS composed of 27 SNPs. **(C)** Median number of individuals who share the same PRS values as a function of number of possible scores. We found that, on average, two individual share the same PRS value in UKBB with ∼10^6^ possible scores, which corresponds to a PRS composed of only 14 SNPs. **(D)** The percentage of identifiable individuals and the median number of individuals who share the same PRS value as a function of number of decimal places reported in effect weights. As we round the effect weight, we see a drastic decrease in the number of identifiable individuals and increase in the median number of individuals who share the same PRS value. This plot is calculated using a PRS model with 48 SNPs for Type-I diabetes (PGS000869 from the PGS catalog [47]) and UKBB individuals. **(E)** Density of subset-sum problem for the genotype recovery using PRS value as a function of number of decimal places reported in the effect weights. As we round the effect weights, genotype recovery becomes a difficult problem to solve, *i*.*e*., the density increases drastically. This plot is calculated using a PRS model with 48 SNPs for Type-I diabetes (PGS000869 from the PGS catalog [47]) and UKBB individuals.

We first asked whether PRS values were unique to individuals. For this, we developed a theoretical estimation of the number of uniquely identifiable individuals in a database based on the database’s size, PRS parameters (effect weights and their precision, i.e., the number of decimal places provided), and allele frequency statistics in the population. As the majority of genotypes behave as independent random variables (SNPs are independent due to the clumping step in PRS modeling [48]), their linear combinations, given a sufficient sample size, follow the normal distribution, as described by the central limit theorem. By incorporating the effect weights and allele frequencies from the population, we could estimate both the mean and standard deviation of this distribution (see Methods). We showed that the total number of possible scores for a given PRS model was determined by either the number of SNPs or by the precision of the effect weights involved in that specific PRS, whichever yielded a smaller value. By using this distribution, we calculated the proportion of scores uniquely attributable to individuals (“uniqueness”), as well as the median number of individuals sharing the same score (“the median anonymity-set size”). The anonymity-set size refers to the number of individuals in a group who are indistinguishable from each other based on the available data. We compared our analytical estimates with the real scores calculated for the 1000 Genomes and UKBB individuals on the PRSs selected from the PGS Catalog for this study. We found that a single PRS model with 20 SNPs in 1000 Genomes or 27 SNPs in UKBB, on average across all weight precisions, sufficed to uniquely identify 95% of individuals (Fig. 5B). Even when the scores overlapped, the same score was often shared by only few individuals. For instance, in a PRS model with 7 SNPs (1000 Genomes) or 14 SNPs (UKBB), the median anonymity-set size was two, meaning that half of the scores were shared by at most two individuals (Fig. 5C). See Supplementary Fig. 4 for the relationship between the number of possible scores and the number of SNPs observed in the actual data (1000 Genomes and UKBB).

Our analytical results slightly overestimate the number of uniquely identifiable individuals, as they do not account for the repetition of effect weights or residual linkage disequilibrium in PRS models, which leads to deviations from the normal distribution and makes some scores more common than predicted. Our analysis pertains to large mixed-cohort databases where the allele distribution conforms to the general population. A disease-specific database might include individuals with a higher rate of risk alleles and hence PRSs at the tail of the distribution, which will decrease identifiability. However, the analysis accurately captures the overall trend, which enables an attacker to estimate the potential success of their de-anonymization attack before accessing a dataset or a data custodian to perform risk assessment. The results presented above pertain to a single trait. The likelihood of two individuals with different genotypes having identical PRSs across multiple traits multiplicatively decreases, as potential overlaps follow the binomial distribution. Thus, several PRSs would provide a robust means of identifying individuals, even in the largest datasets currently available.

This analysis considers a distinct linkage threat model in which an adversary has access to an anonymized genotype database—or to PRS values derived from such data—and seeks to link individuals across datasets by using PRS uniqueness. This access may arise through unauthorized database access, insider misuse, or policy-compliant research use in which identifiers have been removed but genetic data remain available. This access assumption applies only to the PRS-uniqueness analysis presented here and is not required for the genotype-recovery or genealogy-based scenarios analyzed in previous sections. The purpose of this analysis is to quantify identifiability under these assumptions. While statistical uniqueness alone does not guarantee exploitability in real-world settings, it measures the potential for linkage if such access is available.

Next, we investigated a potential strategy to enhance privacy. As the uniqueness of a PRS largely depended on the precision of the effect weights, reducing the number of decimal places in effect weights decreased the uniqueness of each PRS and increased the size of anonymity sets (Fig. 5D). In other words, if PRS values were calculated by using precise weights but the released PRS data were rounded, it became more challenging to match known individuals to anonymous datasets. We further demonstrated this by analyzing PRS solvability and showing that rounding effect weights increased the density of possible solutions (Fig. 5E), thereby making it much harder to predict genotypes from the PRS. Importantly, while rounding effect weights greatly improved privacy, it had almost no impact on the utility of the PRS, as the distributions of PRS values calculated with precise and rounded weights remained closely aligned (Supplementary Fig. 5).

## 3 Discussion

In this study, we identify critical privacy risks associated with the sharing of PRSs. By framing genotype inference from PRSs as the subset-sum problem, we demonstrate that individual genotypes can be accurately predicted from PRSs, achieving a median accuracy of 94.6% across diverse ancestry groups. This capability to reconstruct genetic information from PRSs raises serious privacy concerns, particularly when PRSs are openly shared by research publications or individuals seeking personalized health insights. The implications of these findings are significant for both direct-to-consumer genetic testing companies and large-scale research initiatives.

We further reveal that the risk of inferring genotypes from PRSs is not uniform across populations. Individuals of African and East Asian descent are particularly vulnerable due to differences in allele frequencies between these groups and the European-centric datasets commonly used to derive PRSs. This underscores the need for privacy assessments of PRSs to consider the diverse genetic backgrounds represented in genomic datasets. Additionally, our analysis shows that PRS values can be used to predict health risks beyond the traits originally included in the PRS. This ability to infer supplemental phenotypic information poses challenges for patient consent and data governance frameworks, which often overlook such secondary uses of data. As PRS-based predictions become more integrated into clinical decision-making, a reassessment of current informed consent polices might be needed to ensure that they address these emerging risks.

To address these concerns, we propose a simple yet effective mitigation strategy. Reducing the precision of effect weights in publicly released PRS models can significantly decrease the identifiability of individuals, thus enhancing anonymity within large cohorts while preserving the overall utility of the scores. Concretely, we propose density as a practical tool for selecting PRS disclosure precision. Data holders can progressively round effect weights, compute the resulting density, and determine whether the model enters a regime where likelihood estimation becomes ineffective in identifying correct genotypes among all valid possibilities. Our results show that modest rounding substantially reduces recovery accuracy while preserving overall PRS distributions. Density should be interpreted, however, as a conservative indicator of practical risk rather than a formal guarantee. We also note that while effect-weight rounding substantially reduces privacy risk, high-precision weights are routinely produced and disseminated for reproducibility and reuse. A possible approach for studies is to release two versions of models: one with high-precision weights for reproducibility and another with rounded weights for clinical use.

Finally, integrating privacy-enhancing techniques like differential privacy into the computation and sharing of PRSs might offer a potential safeguard against re-identification, though care must be taken to balance privacy protection with the preservation of PRS utility. Further research is needed to refine these approaches and establish best practices for maintaining privacy while supporting the use of PRSs in personalized medicine. In conclusion, as PRSs gain wider adoption in healthcare and research, our study emphasizes the need for a deeper understanding of the privacy risks that they pose. It is essential to develop robust strategies that facilitate the continued use of genetic data for scientific and clinical progress while safeguarding individual privacy.

### Limitations and future work

In our study, we evaluate the genotype recovery for PRSs based on up to 50 SNPs. This bound reflects a conservative computational constraint rather than a methodological limitation. Genotype recovery is formulated as a subset-sum problem and is therefore NP-hard; as with other problems in this class, practical solvability depends on available computational resources rather than an intrinsic size threshold. PRSs of this scale are, however, not rare—approximately 12% of the models listed in the PGS Catalog that we analyzed contained fewer than 50 variants.

Our analysis of effect-weight rounding focuses on privacy risk mitigation rather than comprehensive evaluation of predictive accuracy. While such assessment would require disease-specific validation and is beyond the scope of this study, Supplementary Fig. 5 indicates that rounded weights preserve the overall PRS distribution across multiple precision levels. We therefore present rounding as a pragmatic risk-reduction strategy, not as a universally optimal clinical solution.

Our results should be interpreted within the specific threat models analyzed. Genotype recovery, genealogy-based re-identification, and PRS-based linkage represent distinct adversarial settings with different access assumptions and operational requirements. We do not claim that the conditions necessary for all three scenarios are simultaneously satisfied in practice. Rather, our goal is to characterize the privacy risks that arise under each plausible access model. In particular, the PRS-uniqueness results demonstrate that individual PRS values can function as strong quasi-identifiers when computed over sufficiently large panels. Whether this translates into a practical linkage attack depends on institutional safeguards, access controls, and governance mechanisms. Taken together, these analyses highlight that PRSs can introduce multiple, conceptually distinct privacy risks.

### Data and Code Availability

This study uses genotypes from the publicly available 1000 Genomes Dataset for genotype recovery and re-identification attacks. It also uses genotypes of the UKBB individuals in the analytical framework to understand the privacy risks associated with the release of PRSs. The code and the data for running a genotype recovery demo, for plotting the figures from this study, and for end-to-end reproducibility of the experiments can be found at https://github.com/G2Lab/prs-privacy.

## 4 Methods

### 4.1 Overview of datasets

In all our experiments, we use publicly available genomic datasets, specifically the phase 3 release of the 1000 Genomes project (2,535 samples) [49] for privacy attacks and the imputed version of the UKBB dataset (487,409 samples) for uniqueness and anonymity estimations [50]. The 1000 Genomes dataset is comprised of 2,467 unrelated individuals, and 25 and 9 pairs of first- and second-degree relatives, respectively. We use the relatedness information for de-anonymization.

We obtain PRS metadata from The Polygenic Score (PGS) Catalog [29], an open database of published polygenic scores. The catalog provides metadata for a PRS as a scoring file that specifies the trait, the information source, and the details about the associated variants. We calculate the target PRSs and population allele statistics from the datasets’ genotype data, and we use the ancestry and kinship information from the datasets’ metadata where appropriate. We do not attempt to infer any unknown attributes or identities of the individuals.

### 4.2 Preprocessing of PRSs from the PGS catalog

We collected all available PRSs with their metadata for any PRS with *N <* 50 SNPs from the PGS catalog. We filtered them by first removing the scoring files that unfairly simplify our task: PRS with a single variant *N* = 1 and with a high difference between the precision across its effect weights. We then removed the PRSs with (1) missing or incorrectly specified loci; (2) the effect alleles that did not match either the reference or alternative allele in 1000 Genomes data; (3) SNPs on the X or Y chromosomes, and (3) density *d >* 2.5. In total, we ended up with 298 PRSs spanning 4,821 total loci, out of which 2,654 were unique (see Supplementary Table 1 for the filtering details).

### 4.3 Framing genotype prediction as the subset-sum problem

A polygenic risk score is a linear combination of the genotypes *g* ∈ {0, 1, 2} of the SNPs associated with a disease and the corresponding effect weights *β*. The score is typically normalized by the number of SNPs *N* and the ploidy *P* (*P* = 2 for diploid cells):

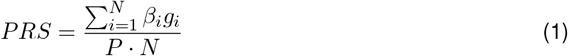

The effect weights are real numbers that represent the contribution of each genetic variant (often single nucleotide polymorphisms, or SNPs) to the overall risk of developing a particular disease. Studies release effect weights with varying degree of precision (the number of decimal places).

We posit that the task of finding the genotypes that yield a given PRS value is a variation of the subset-sum problem, where one attempts to find a subset of a number set that sums up to a target value. This problem is known to be NP-hard (Non-deterministic Polynomial-time hard) [33], and, hence, computationally difficult to solve. However, there are algorithms that can solve this problem efficiently when certain conditions are met regarding the properties of the numbers involved [36, 34, 51, 52], and these conditions can be met in the context of PRS. Thus, we reduce the genotype recovery task to an instance of the subset-sum problem by defining the effect weights as the number set and the PRS value as the target sum. The number of times each effect weight (0, 1, or 2) is used in the solution determines the genotype for each corresponding SNP.

#### Solvable PRS

Solving an NP-hard problem even with an efficient algorithm is computationally intensive process when the problem’s size is large. Hence, it is important to be able to estimate the feasibility of genotype recovery before attempting to solve it. The subset-sum problem’s difficulty is often measured by using a concept called *density*. This density is defined as the ratio between the number of elements in the set and the bit length (or the number of binary digits) of the largest number in that set [34]. The idea is that (1) the more numbers a set has, the more possible subsets can be formed, and (2) the shorter the bit length of the numbers, the lower the precision and the fewer possible sums that can be achieved. Naturally, problems with a higher density are more difficult to solve [35] because multiple subsets can yield the same sum.

In contrast to the classic subset-sum problem where the numbers are a set of integers, PRS effect weights are real values and, hence, not all of them can be precisely represented with bits. In addition, genotypes can be one of the three possible values, so we need the ternary bit representation of the largest weight. We adapted the bit precision estimation to the PRS context by calculating base-three logarithm of the number of decimal places in the largest weight. The number of decimal places must be normalized across all the weights, i.e., if any of other weights are “longer”, we append zeros to the largest weight to match the decimal length. The density is then calculated as the ratio between the total number of effect weights and the largest weight’s ternary-bit representation length:

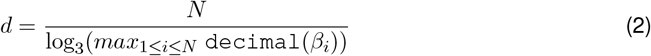

Prior work has primarily focused on low-density problems [34, 52, 51, 35], i.e., *d <* 1, as most studies considered applications to cryptography, where problems have a single solution. Given our approach of combining dynamic programming with likelihood estimation (see below), we hypothesized that we would be able to solve denser problem instances. To determine a feasible threshold, we selected 268 PRS models of up 30 SNPs and performed genotype recovery for 50 individuals from the 1000 Genomes dataset for different densities by gradually rounding the effect weights of each model. Our results showed (Supplementary Fig. 6) that for density *d >* 2.5, the recovery advantage over predicting the major genotype for each SNP becomes small, which was consistent with the performance of the best-known algorithms [53, 54]. We note, however, that density should be used only as a conservative indicator of practical solvability and not as a guarantee of uniqueness or correctness of recovered genotypes.

Resource and density constraints ultimately limit the size of a PRS instance that our algorithm can handle. The computational and memory requirements grow superlinearly with the number of SNPs *N* and their precision. In order to analyze the requirements in practice, we experimentally measured the CPU time and peak memory usage of genotype recovery across the previously selected 298 PRS models (see Supplementary Figure 7). In this work, we focus on PRS instances with *N <* 50, but we estimate that an adversary with sufficient computational resources could extend this limit to around *N <* 80. Most studies choose decimal-places weight precision of less than 15, and, hence, the problem becomes too dense to be easily solvable after 80 SNPs.

#### Our Dynamic Programming Approach

We chose to use dynamic programming to solve the genotype recovery posed as the subset-sum problem as it had the best worst-case performance [36]. Other efficient algorithms [34, 51, 52] were either based on heuristics, and hence did not guarantee finding the best solution, or were applicable only to low-density problems. We combine classic dynamic programming with several optimizations techniques [37] to reduce the computational costs. We first split the effect weights into two equal-size subsets and build a sums table for each. The building process consists of iteratively adding each effect weight (either once or twice to account for one or two effect alleles at the SNP) to the existing weight sums in the table. During each addition, a pointer is recorded to track the last weight added for each sum. This method enables us to construct all the weight sequences that produce each sum in the table. Once the tables are constructed, we use the meet-in-the-middle technique of iterating over all the sums in the first table and checking if the difference between the target and a sum matches any sum in the second table. When we find a match, we backtrack through the tables to find the corresponding weight sequences and translate them into the genotypes, whose linear combination sums up to the target PRS.

The meet-in-the-middle technique enables us to reduce the memory complexity from *O*(3^*N*^) of the direct approach to *O*(3^*N/*2^). In order to reduce the costs further, we maintain upper- and lower-bounds on the intermediate sums that still can lead to the target score, given the remaining weights. If a sum is either larger than the target or the remaining weights do not suffice to reach the target, we do not add this sum to the table. If some of the weights are negative, we extend the bounds in both tables, accordingly. Note that the presence of negative weights makes solving the problem more resource-intensive [55]. Finally, in order to reduce the size of tables and, hence, the computational requirements, we divide the search into two stages. First, we build the tables with rounded-down weights and find solution candidates at rounded-down precision. Then, we calculate the score of each candidate at the full precision and select the solutions that match the target.

#### Choosing the best solution

Dynamic programming enables us to find *all* valid genotype combinations that result in a given PRS. We still need a way to select the “best” one among them. The insight that helps us is that variations in the human genome are not uniformly distributed—some genotypes are more likely than others. Hence, we can quantify the likelihood of a genotype for a given variant by its frequency in the population. To simplify the analysis, we consider the SNPs in a PRS model statistically independent from each other and measure the the likelihood of a solution by combining the likelihoods of the individual SNP genotypes. Specifically, we calculate the sum of the log-probability for all the solution genotypes based on the allele frequency in the population that the target individual belongs to (AF) and select the solution with the highest sum as the best one. This approach consistently places the true solution in the top ranks of all valid solutions for majority of individuals. In practice, PRS models might contain correlated variants due to residual Linkage Disequilibrium. If an attacker identifies such highly correlated variants, they can use it to their advantage by boosting the likelihood of the more probable SNP combination or even avoid adding the unlikely combination to the sums tables, and hence increasing the prediction accuracy or reducing the computational overhead.

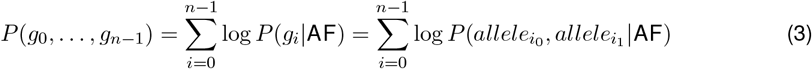

As the number of SNPs included in a PRS is large, the number of possible solutions grows rapidly, making it impractical to keep track of or to store all the potential solutions. We address this issue with an algorithm that, instead of storing all possible paths, keeps a reference only to the path with the highest likelihood for each sum in the table. The process involves three steps: (1) calculating the likelihood of a solution where all genotypes consist of non-effect alleles, (2) updating the likelihood for each new sum by adjusting the log-probability—subtracting that of the non-effect alleles and adding that of the effect alleles—to the highest likelihood from the previous sum, and (3) keeping a reference to the last added effect weight that maximizes the likelihood for each sum. During backtracking, we trace only the path with the highest likelihood at each step, ultimately identifying the solution with the greatest likelihood. The complete algorithm for recovering genotypes from a PRS is outlined in Supplementary Algorithm 1.

#### PRS chaining and Self-repair

In a realistic scenario, an attacker might have access to multiple PRS values from a single individual. Some PRSs share overlapping SNPs, as the same genetic mutation can influence multiple traits. The attacker could exploit this overlap to infer genotypes for these SNPs. The strategy is to first predict genotypes for the PRSs with fewer SNPs, as they can be computed quicker. Hence, we first solve the PRS with the smallest set of SNPs. For each subsequent PRS that includes more SNPs, we retain the genotypes of overlapping SNPs from the previous solution. This approach gradually reduces the possible solution space as the number of SNPs increases, thus making the computations more efficient.

When inferring genotypes from a single PRS, the attacker lacks a direct way of confirming whether their highest-likelihood solution contains the correct genotypes. However, access to multiple PRSs from the same individual provides a way to cross-check solutions and to increase recovery accuracy. If the attacker cannot find a solution for a PRS using previously inferred genotypes, it likely indicates that some of the earlier predictions were incorrect—for example, the correct solution might have had the second highest likelihood. To address this, we propose a “self-repair” strategy. When a PRS fails to yield a solution with the genotypes predicted from a smaller PRS, the attacker attempts to predict the genotypes without using the genotypes predicted from a smaller PRS and generates a sorted list of potential solutions. The attacker then tests each solution in the list, checking whether it enables successful prediction of genotypes from all previously processed PRSs. If a solution validates the genotype prediction for earlier PRSs, the attacker updates their genotypes with this refined solution and proceeds to the next PRS. As genotype solutions are ranked by using population-based likelihoods, incorrect early solutions are therefore unlikely to remain top-ranked as more PRSs are incorporated. This iterative strategy gradually fixes early prediction errors.

Similarly to exploiting residual LD in a single PRS model, our approach can, in theory, be extended to correlated variants across multiple models. In addition to the overlapping SNPs, the attacker can also fill the correlated SNPs of larger PRSs and thus further reduce the solution search space.

### 4.4 Linking predicted genotypes to databases

Genealogy services enable users to upload their genotype information to the database and search for their potential relatives among other users. The attacker can upload the recovered genotypes of the individual to a genealogy database and discover the genetic matches. The exact algorithms that these services use are not disclosed but prior studies [41, 42] suggest that they search for long genomic segments with no mismatching homozygous sites between pairs of individuals. Here, we emulate a genealogy search with the individuals from the 1000 Genomes dataset, by using popular algorithms for kinship inference from SNPs, namely KING-robust [43] and GCTA [44].

We first created a database by using the 2,467 unrelated individuals, 25 pairs of first-degree (parent-child, siblings) and 9 pairs of second-degree relatives (grandparent-grandchild, half-siblings, etc) from the 1000 Genomes data. After predicting ∼2,600 genotypes with ∼5% error for each individual, we linked them to the emulated database that contained the full set of genotypes for each individual. We used PLINK [56, 57] to compute a matrix with pairwise kinship coefficients for both KING-robust [43] and GCTA [44] algorithms. We then used the relationship inference criteria of 0.354 for self-identification, 0.177 for first-degree relatives, and 0.089 for second-degree relatives [43, Table 1]. Having the coefficients for all the pairs enabled us to calculate the precision and recall of linking each individual to themselves and their relatives, if the latter were present, and to measure the distribution of the coefficients for each category.

#### KING-robust

KING-robust is an algorithm for pairwise relationship inference in the presence of population substructure. The algorithm calculates the kinship coefficient that estimates the probability of two randomly sampled alleles from two individuals being identical by descent (IBD). The high-level idea is that having different homozygous alleles at the same locus reduces the IBD probability. The kinship coefficient is calculated as a relation between the number of loci where two individuals both have heterozygous alleles (*N*_*Aa,Aa*_ for reference allele *A* and alternative allele *a*), have different homozygous alleles (*N*_*AA,aa*_), and the total number of heterozygous alleles (*N*_*Aa*_):

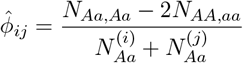

#### GCTA

GCTA is also an algorithm for pairwise estimation of genetic relationship. It measures kinship in terms of how much the genotypes that two individuals have deviate from the expected counts in the population. As it relies on the allele frequency in the population, it is less robust than KING-robust for individuals with different ancestry. Given the number *x*_*k*_ of the reference alleles that an individual has and the frequency *p*_*k*_ of the reference allele in the population for the *k*^*th*^ SNP, the kinship between individuals *i* and *j* with *N* SNPs is calculated as follows:

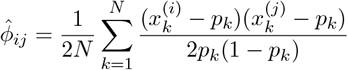

### 4.5 PRS as an identifier to link known individuals to anonymized databases

Given access to an anonymized genomic dataset and the PRS of a known individual, the attacker can compute the PRS for each sample in the dataset. If one matches the target PRS, the attacker could infer the individual’s genotypes or associated phenotypes without the need for computationally intensive genotype recovery. This method relies, however, on the assumption that a PRS uniquely identifies the individual. In practice, different genotype combinations can yield the same score, and multiple samples may share an identical set of genotypes for the SNPs in the PRS model.

Here, we demonstrate how an attacker can assess the uniqueness of a given PRS within a population, thereby estimating their confidence that a potential match corresponds to the individual of interest. Intuitively, the uniqueness of a PRS is influenced by factors such as the number of included SNPs, the precision of effect weights (i.e., the number of decimal places), allele frequencies in the population, and the size of the target dataset or population.

#### PRS as the normal distribution

As genotypes in PRS models are supposed to be independent due to LD clumping, the central limit theorem implies that their linear combination will approximate a normal distribution, given a sufficiently large sample size and a sufficient number of variants in the PRS. This normal distribution is characterized by its mean and standard deviation; therefore, estimating these parameters enables us to determine the probability density associated with any given PRS.

Under the assumption that PRS variants in individuals from mixed non-disease cohorts adhere to the Hardy-Weinberg equilibrium, we can estimate the expected contribution of variant *i* to the PRS based on the population allele frequency *p*_*i*_ and the effect weight *β*_*i*_:

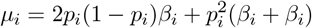

Then, the expected mean *µ* and standard deviation *σ* of a PRS, given the ploidy *P* and the number of variants *N*, become

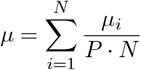

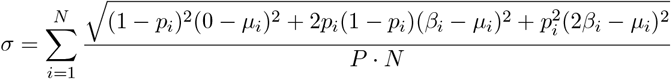

For a PRS to approximate a normal distribution, its effect weights must be densely and evenly distributed. If one weight, for instance, is orders of magnitude larger than the others, it can skew the distribution, potentially creating a trimodal shape. In this study, we observed that most PRSs exhibited the normal distribution. This is likely because associations with polygenic diseases or traits are typically influenced by multiple variants rather than being dominated by any single one.

#### Analytical estimation of score uniqueness

Having estimated mean *µ* and deviation *σ*, we can use the probability density function of the normal distribution to calculate the probability of a score *x* being in the range [*x* − Δ, *x* + Δ] as

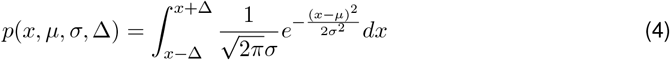

In order to assess score uniqueness, however, we need the probability of each specific discrete score rather than a continuous range. We reconcile this requirement with the continuousness of the normal distribution by defining the probability of a discrete score *x*_*i*_ as the probability of the score being in the range [*x*_*i*_ − Δ, *x*_*i*_ + Δ], where 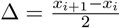.

The total number of possible scores and, hence, distance Δ are determined by either the total number of possible combinations of the effect weights or their decimal precision. Given a PRS with *N* variants and *B* distinct weights {*β*_0_, …, *β*_*B*−1_}, we can calculate the number of possible weight combinations as 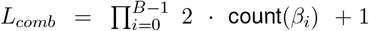, where function count returns the number of variants that share the same effect weight. For example, *L*_*comb*_ = 3 · 3 · 5 = 45 for a PRS with four variants, two out of which share the same effect weight. We approximate the range of possible scores due to the effect weights’ decimal precision for a PRS with *N* variants as:

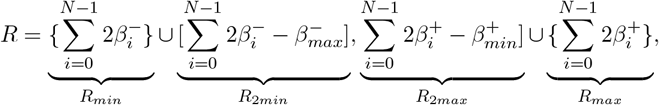

where *β*^+^ and *β*^−^ are positive and negative effect weights, respectively. We exclude the ranges [*R*_*min*_, *R*_2*min*_] and [*R*_2*max*_, *R*_*max*_] as an optimization because we know for certain that a PRS cannot take any value from these segments.

The number of possible scores due to the decimal precision becomes

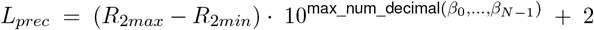

We then can set *L*_*scores*_ = min(*L*_*comb*_, *L*_*prec*_) and calculate 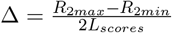.

Given the score probability defined in Eq. (4) and calculated Δ, we can estimate the probability of a single sample in a dataset of size *M* to have score *x*_*i*_, i.e., being uniquely identifiable, by using the probability mass function of the binomial distribution. In essence, we calculate the probability that *M* trials of an event with probability *p*_*i*_ results in only one successful outcome.

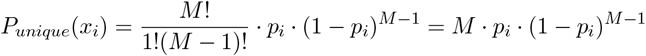

To calculate the expected total number of unique scores, we sum the probabilities of each score *x*_*i*_ being unique:

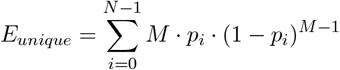

To estimate the median number of samples in the dataset with the same score, we calculate the expected number of samples for each score *x*_*i*_ ∈ *R* as *E*(*x*_*i*_) = *M* · *p*_*i*_, store all the results with *E*(*x*_*i*_) ≥ 1 in an array, then sort the array and return the middle element.

Finally, we partially decrease the computational overhead of the analysis by calculating Eq. (4) with the cumulative distribution function (CDF) of the normal distribution:

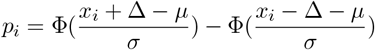

where Φ is the CDF of the normal distribution.

## A Supplementary Information

### A.1 Supplementary Figures

**Supplementary Figure 1.**
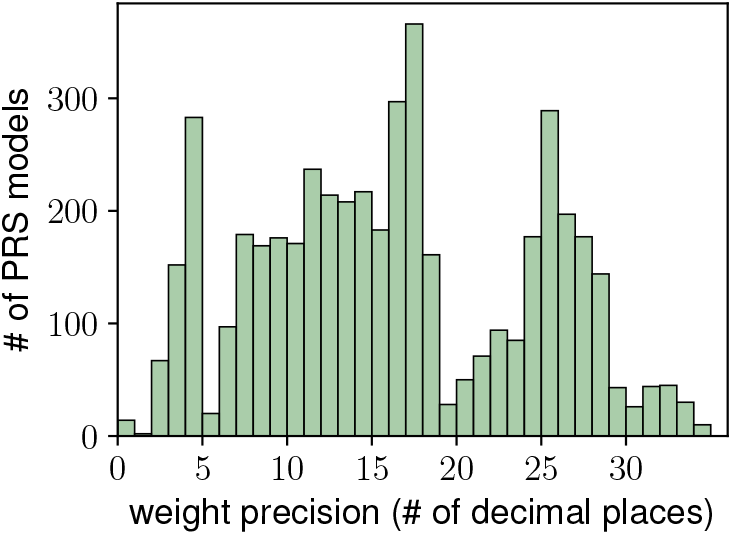
The distribution of effect-weight precision across all PRS models published in the PGS Catalog [29]. The median precision is 15 digits.

**Supplementary Table 1.**
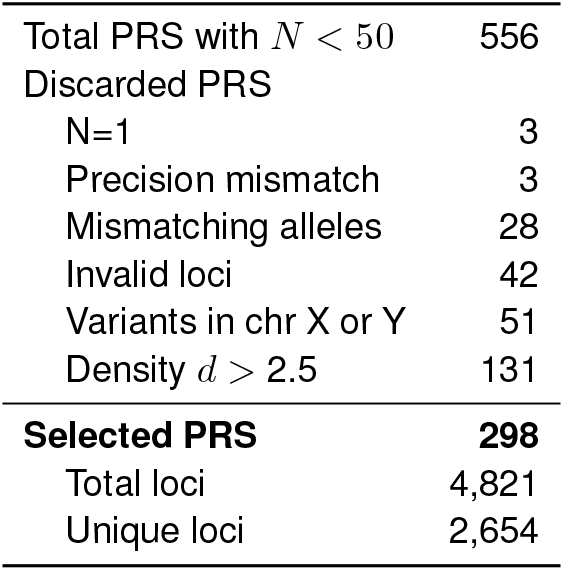
The PGS Catalog selection for the experimental evaluation.

**Supplementary Figure 2.**
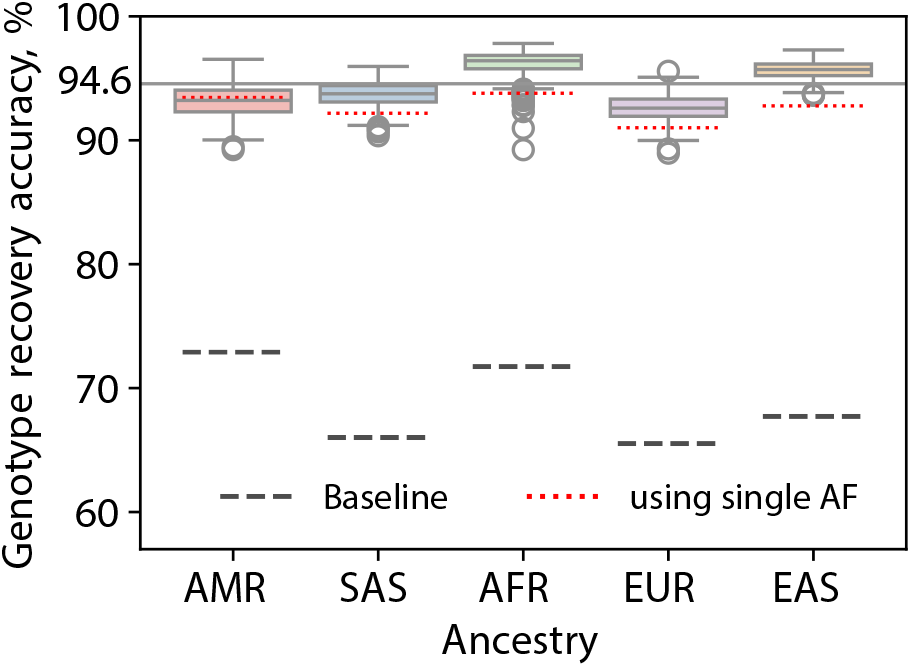
Genotype recovery accuracy. Genotype recovery accuracy across different populations: African (AFR), Admixed American (AMR), East Asian (EAS), European (EUR), and South Asian (SAS). Baseline represents the recovery accuracy when we predict the major allele for each SNP for every individual. The red lines are the accuracy when we use a single common allele-frequency statistic for all the populations in our likelihood estimations.

**Supplementary Figure 3.**
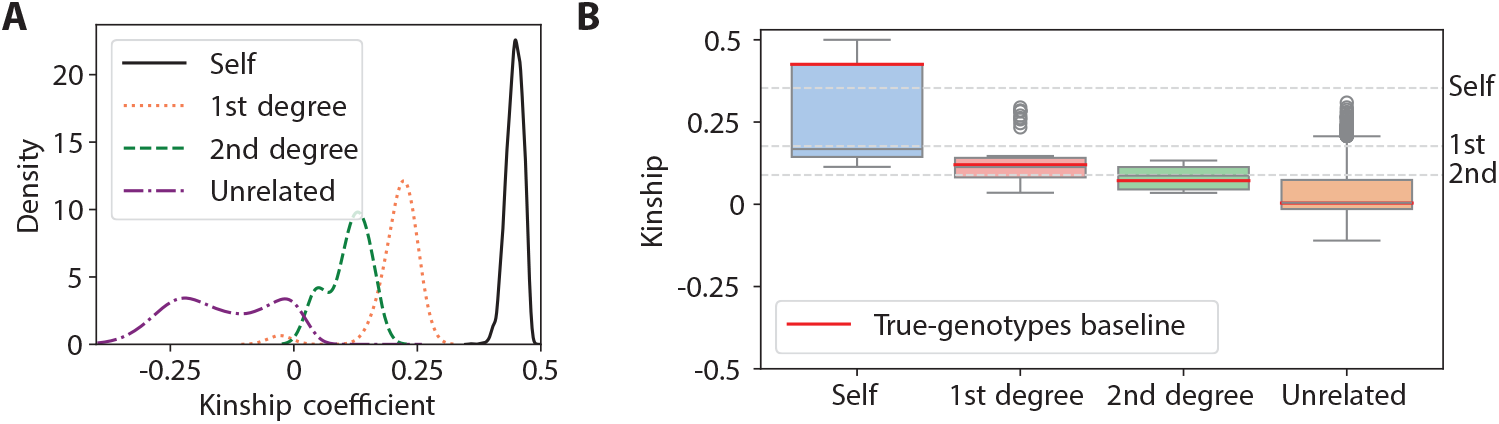
Kinship analysis. **A** The distribution kinship coefficients determined with our predicted SNPs by using the KING algorithm. Compare this figure with the distribution of kinship estimates on 5,000 SNPs from the KING paper [43, Figure 1C]; **B** The kinship coefficients for the target individuals and their first- and second-degree relatives calculated by using GCTA algorithm.

**Supplementary Figure 4.**
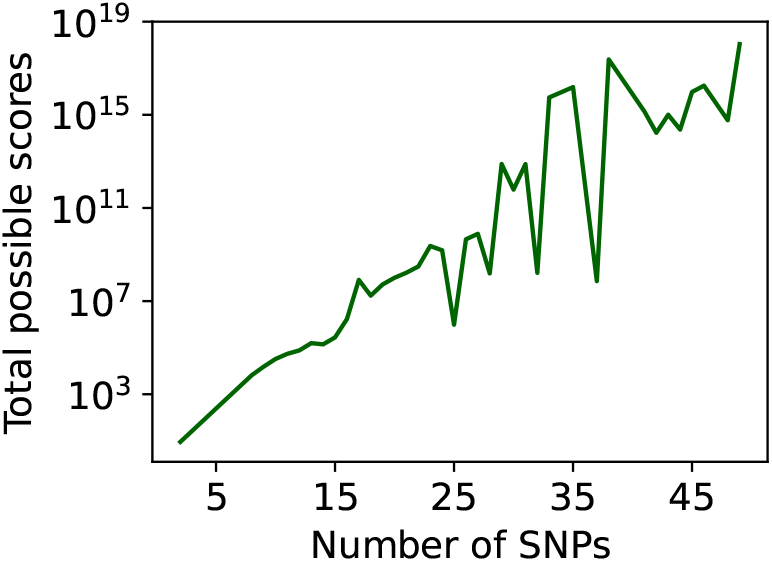
The total number of possible PRS values in a database vs. the number of involved SNPs. The total number of possible scores per number of SNPs are calculated by using the UKBB data and by taking the median values for different effect weight precisions (i.e., the number of decimal places)

**Supplementary Figure 5.**
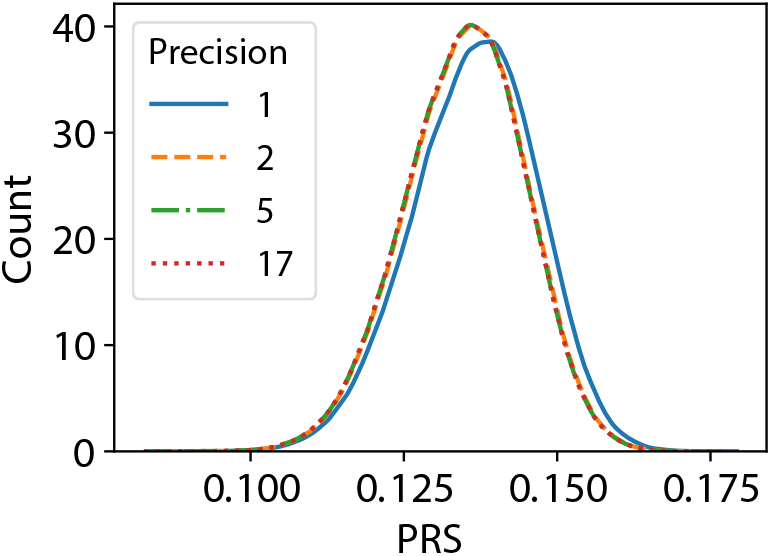
PRS vs. effect weight precision. The PRS distribution for varying levels of precision in effect weights. PRS values are shown with different levels of precision: 1, 2, 5, and 17 decimal places.

**Supplementary Figure 6.**
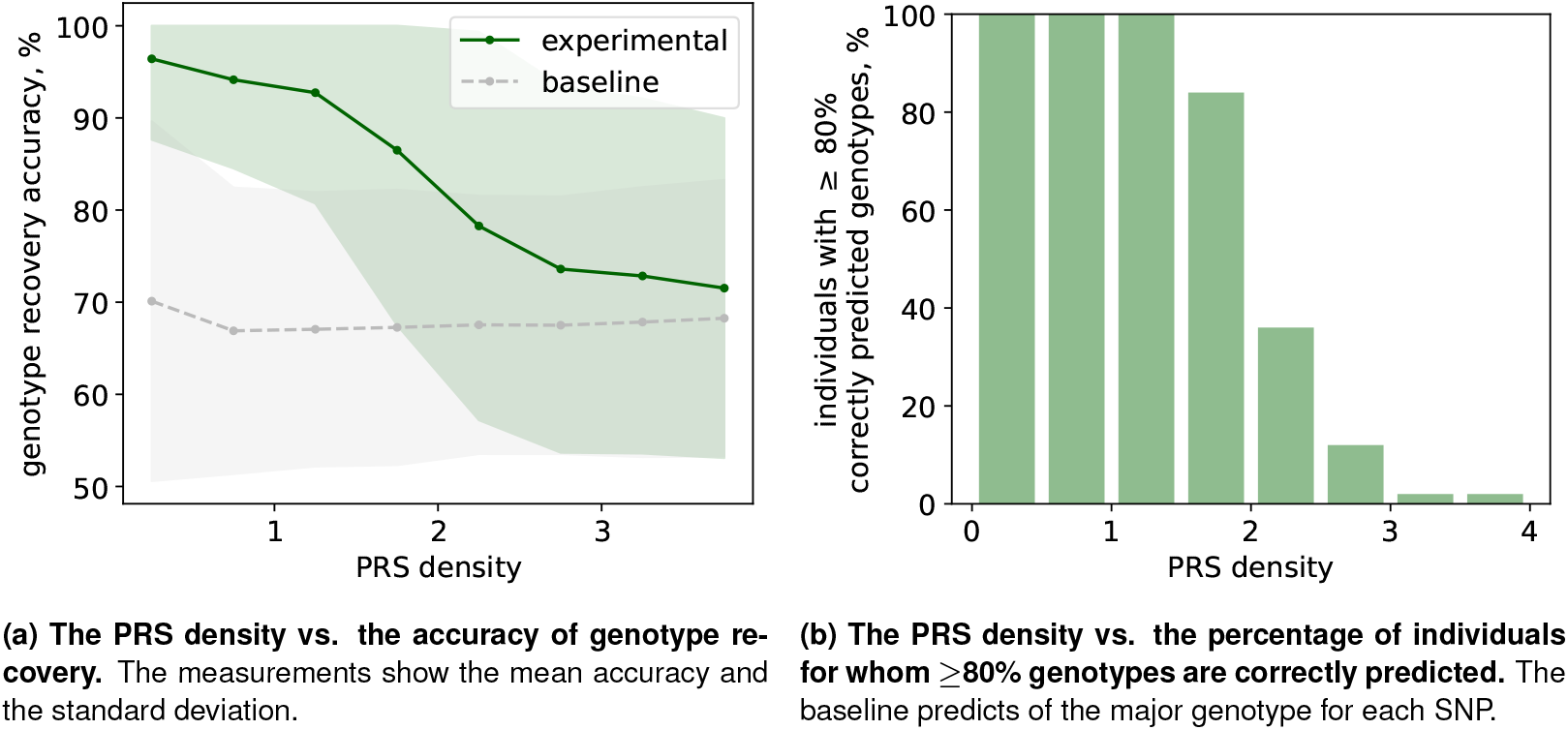
The accuracy of genotype recovery for 268 PRS models of up to 30 SNPs (Supp. Table 1) and 50 individuals from the 1000 Genomes dataset. The density of each model is gradually increased by rounding the effect weights one decimal place at a time.

**Supplementary Figure 7.**
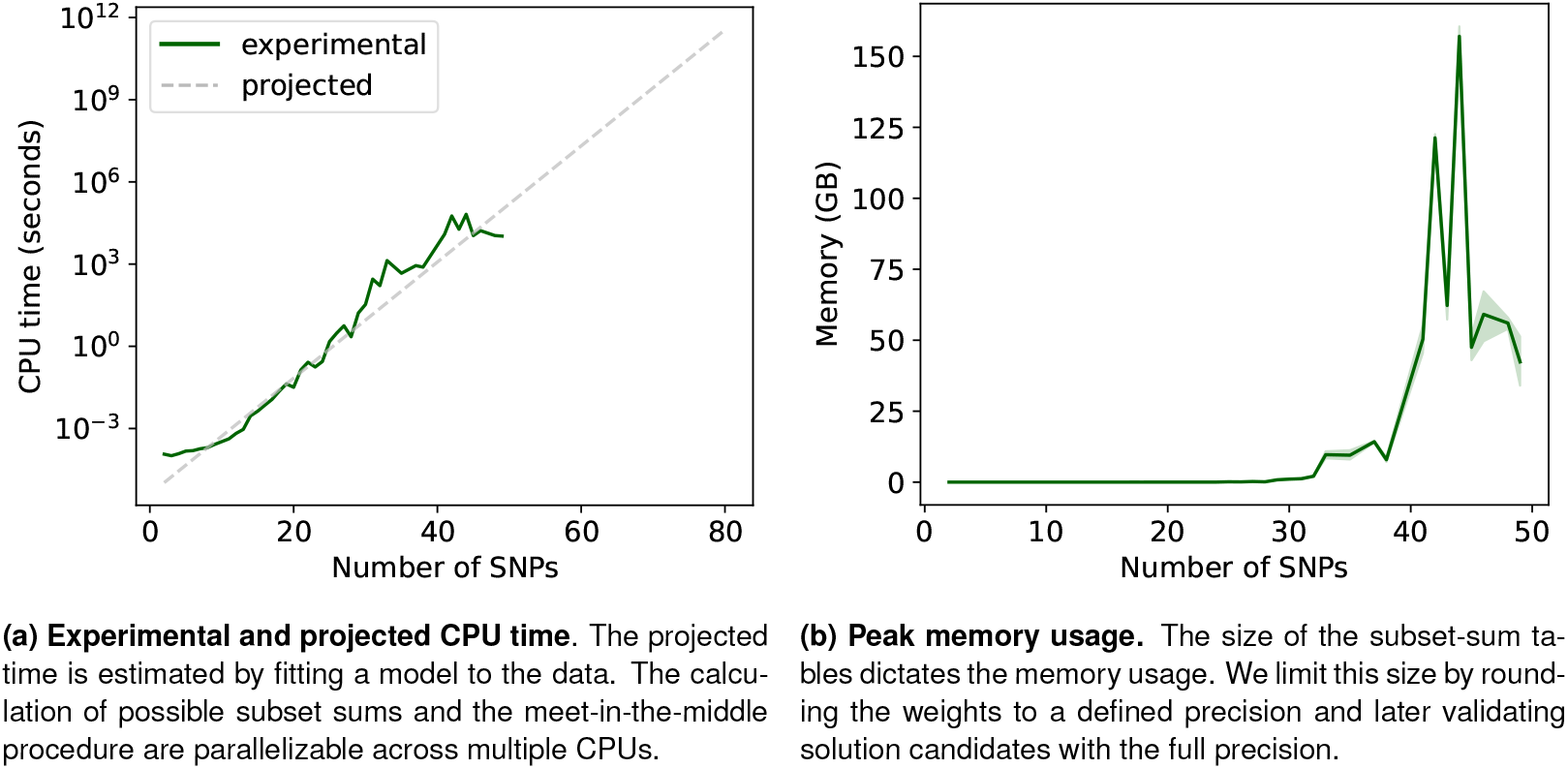
The experimental measurement of the resource usage during the genotype recovery averaged across 298 PRS models (Supp. Table 1) and 50 individuals from the 1000 Genomes dataset.

### A.2 Supplementary Algorithms

#### Algorithm 1 GenotypeRecovery

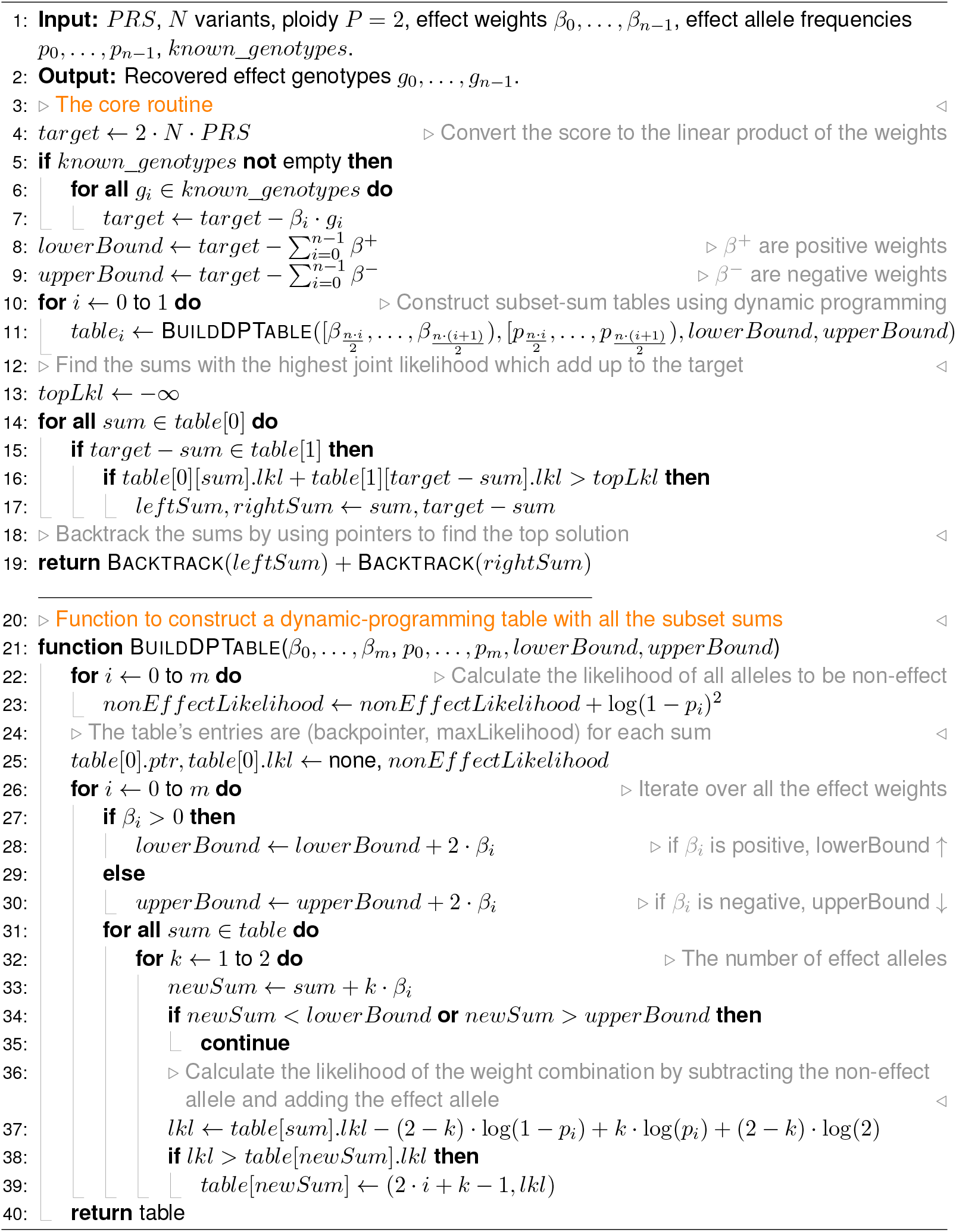

